# Identification of novel Alzheimer’s disease genes co-expressed with *TREM2*

**DOI:** 10.1101/2020.11.13.381640

**Authors:** Joseph S. Reddy, Mariet Allen, Xue Wang, Joanna M. Biernacka, Brandon J. Coombes, Gregory D. Jenkins, Jason P. Sinnwell, Minerva M. Carrasquillo, Cyril P. Pottier, Yingxue Ren, Vivekananda Sarangi, Curtis S. Younkin, Yan W. Asmann, Owen A. Ross, Rosa Rademakers, Todd E. Golde, Nilüfer Ertekin-Taner, Steven G. Younkin

## Abstract

By analyzing whole-exome data from the Alzheimer’s disease sequencing project (ADSP), we identify a set of 4 genes that show highly significant association with Alzheimer’s disease (AD). These genes were identified within a human *TREM2* co-expression network using a novel approach wherein prioritized polygenic score analyses were performed sequentially to identify significant polygenic components. Two of the 4 genes (*TREM2*, *RIN3*) have previously been linked to AD and two (*ATP8B*4, *IL17RA*) are novel. Like *TREM2*, the 2 novel AD genes are selectively expressed in human microglial cells. The most significant variants in *ATP8B4* and *IL17RA* are non-synonymous variants with strong effects comparable to the APOE ε4 and ε2 alleles. These protein-altering variants will provide unique opportunities to further explore the biological role of microglial cells in AD and help inform future immune modulatory therapeutic development for AD.

## Background

In Alzheimer’s disease (AD), amyloid β protein (Aβ) oligomerizes and deposits as insoluble amyloid fibrils in senile plaques which reside in the brain for a long prodromal period during which tau protein is deposited in neurofibrillary tangles and mild cognitive impairment occurs followed by dementia^1,2^. It has long been known that the amyloid deposition which occurs in AD is associated with activation of the innate immune system in the brain^1,3,4^. It is well-established that heterozygous *TREM2* variants strongly increase risk of AD^5,6^. *TREM*2 is selectively expressed in human microglial cells^7^, and *TREM2* expression is increased both in the brains of AD patients^8,9^ and in mouse models of amyloid deposition^1,10^. Recent studies of mouse models indicate that *TREM2* plays an important role in regulating the response of the immune system to Aβ and tau pathologies^11–14^. A weighted gene co-expression network analysis of amyloid-bearing mice has shown that *TREM2* is a hub gene in an AD co-expression network activated by amyloid^15^. In another study *TYROBP*, the signaling partner of *TREM2*, was found to be a key regulator in a human immune gene regulatory network relevant to AD pathology^16^. Thus, in principle, therapies which effectively target *TREM2* and other genes in its co-expression network might halt or slow progression to dementia in cognitively normal subjects with amyloid deposition by modulating the immune response that occurs when amyloid is deposited. In an effort to identify novel AD genes co-expressed with *TREM2*, we employed a novel approach based on polygenic scores (PGS) and sequence kernel association testing (Fig.1) to explore 234 genes in a *TREM2*-containing co-expression network (CEN_TREM2_). The genes forming CEN_TREM2_ were identified using weighted gene co-expression network analysis^17^ (WGCNA) of RNAseq data^18^ from postmortem temporal cortex of 80 AD and 76 control brains (Methods).

**Figure 1.**
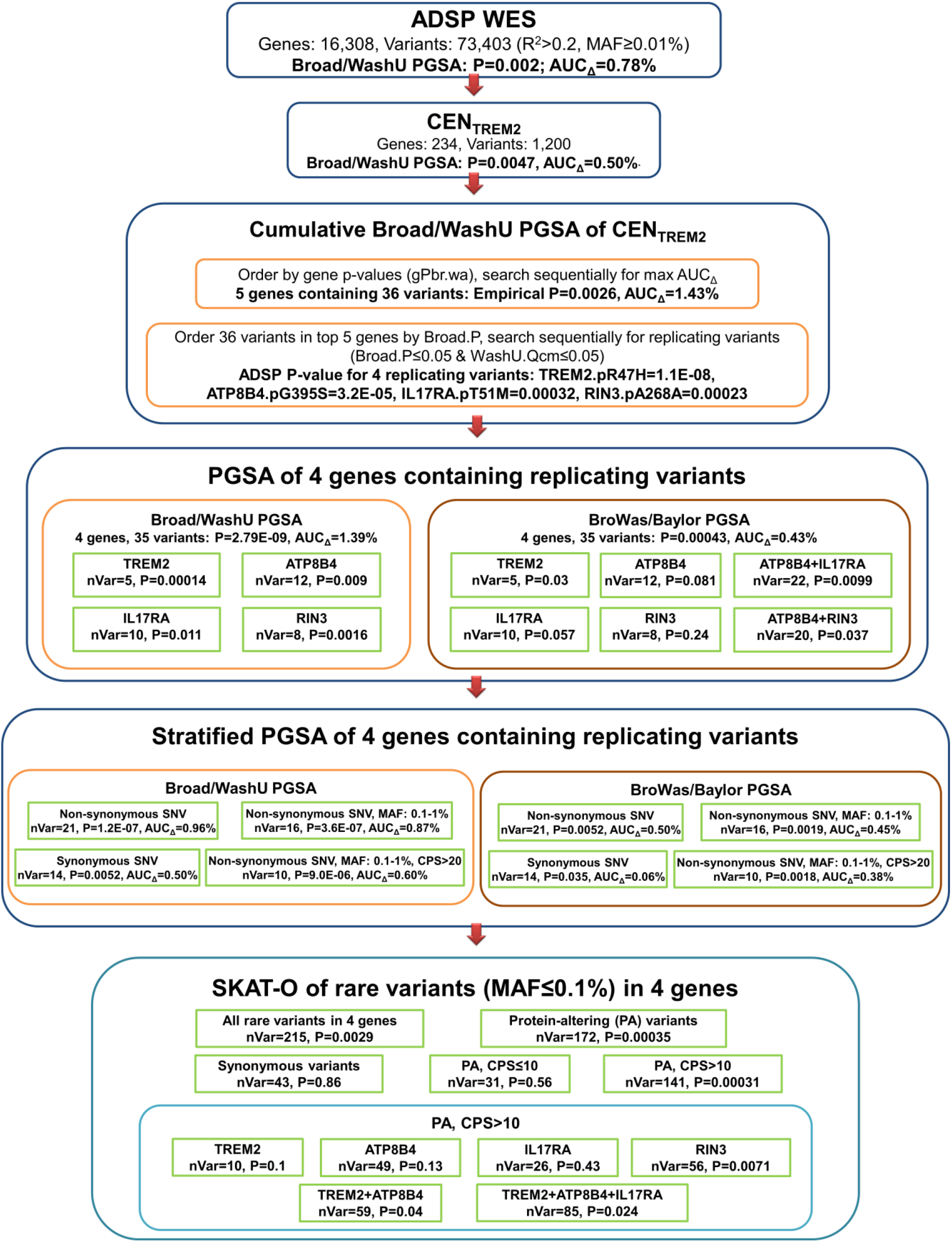
Analytic flow diagram. The ADSP WES and then the 234 genes in a co-expression network containing TREM2 (CENTREM2) were analyzed in sequential analytic steps. Broad/WashU PGSA was employed to analyze pruned variants (r^2^ < 0.2) with MAF = 0.1% thereby identifying a polygenic component (g4v35) comprised of 4 genes with 35 variants wherein each gene showed significant association with AD in the WashU (test) data and had a strongly associating variant that showed significant association both in the Broad (discovery) and WashU (test) data. BroWas/Baylor PGSA was employed for follow-up analysis of g4v35 in independent samples sequenced at Baylor. SKAT-O was used for follow-up analysis of 215 variants with MAF ≤ 0.1% in the 4 genes. AUC_Δ_ is the improvement in AUC that occurred when polygenic scores were added to a covariates only model that included *APOE* ε4 dose, *APOE* ε2 dose, and three principal component vectors. nVar is the number of variants in the genes or polygenic components analyzed. Stratified analyses were performed after stratification on non-synonymous SNVs (nsyn), synonymous SNVs, minor allele frequency (MAF), protein-altering variants (indels + stopgain + nsyn), or CADD PHRED-scaled scores (CPS)..

## Results

### Single Variant Analysis

The ADSP WES dataset^19^ was generated by sequencing a total of 10,929 (dbGaP Study Accession: phs000572.v4.p2) subjects at three large scale sequencing and analysis centers (LSACs). In this dataset, samples from 9904 subjects passed stringent quality control (QC) as fully described in the Methods section. Single variant analysis was performed on 102,828 exonic variants which passed QC and had a minor allele count of twenty or more and a MAF of 0.1% or more. Analysis of these variants, performed by logistic regression using an additive model with sex*, APOE* ε4 dose, *APOE* ε2 dose, LSACs, and three principal component vectors as covariates, yielded five variants with study-wide significance (4.86E-07). As a final QC measure, multinomial regression was performed to assess heterogeneity in minor allele frequency across the three LSACs, and 677 variants (0.66%) with study-wide P_LSAC_ values ≤ 4.86E-07 were removed. Four of the five variants which associated with AD at study-wide significance were among those removed. All four variants showed striking heterogeneity across the LSACs with P_LSAC_ values less than 1.0E-34 even though they passed all other standard quality control measures. Only rs75932628 encoding *TREM2* p.R47H, which is known to associate with Alzheimer’s disease^5,6^, showed study-wide significance after this final QC measure. Supplementary Fig. 1 shows a quantile-quantile (Q-Q) plot comparing single variant results before and after removal of variants with P_LSAC_ values ≤ 4.86E-07. Supplementary Table 1 shows results for all single variants tested and includes P_LSAC_ values, MAF information, and annotation for each variant.

### PGSA of all ADSP and CEN_TREM2_ variants

Of the 9904 post-QC samples, 4452 (45%) were sequenced at the Broad Institute, 3260 (33%) at Washington University in St. Louis, and 2217 (22%) at Baylor University (Supplementary Table 2). To avoid any signal from *APOE*, we removed 122 variants in linkage disequilibrium with *APOE*. To evaluate all remaining variants with MAF > 0.1% for association with AD, we performed polygenic score analysis using Broad data for discovery and WashU data for testing. Baylor data were reserved for follow-up analysis. Variants were clumped (r^2^ < 0.2) using the Broad data to prioritize significant variants from the single variant analysis. We then used the variant effect size estimates based on the Broad data to create polygenic scores (PGS) for AD within the WashU data set. The PGS in WashU were tested for association with AD by logistic regression with sex*, APOE* ε4 dose, *APOE* ε2 dose, and three principal component vectors as covariates. The PGS derived from the Broad sample showed significant association in the WashU dataset (73,403 variants, 16,308 genes, P_PGS_ = 2.05E-03) as did the PGS restricted to variants in CEN_TREM2_ (1200 variants, 234 genes, P_PGS_ = 4.73E-03) and the PGS with variants in CEN_TREM2_ removed (72,203 variants, 16,074 genes, P_PGS_ = 4.94E-03). Consistent with these results, Q-Q plots of the P_ADSP_-values for (i) all ADSP variants, (ii) CEN_TREM2_ variants, and (iii) the ADSP variants remaining after removal of CEN_TREM2_, all deviated from the distribution expected under the null hypothesis of no association with AD (Fig. 2A, B).

**Figure 2.**
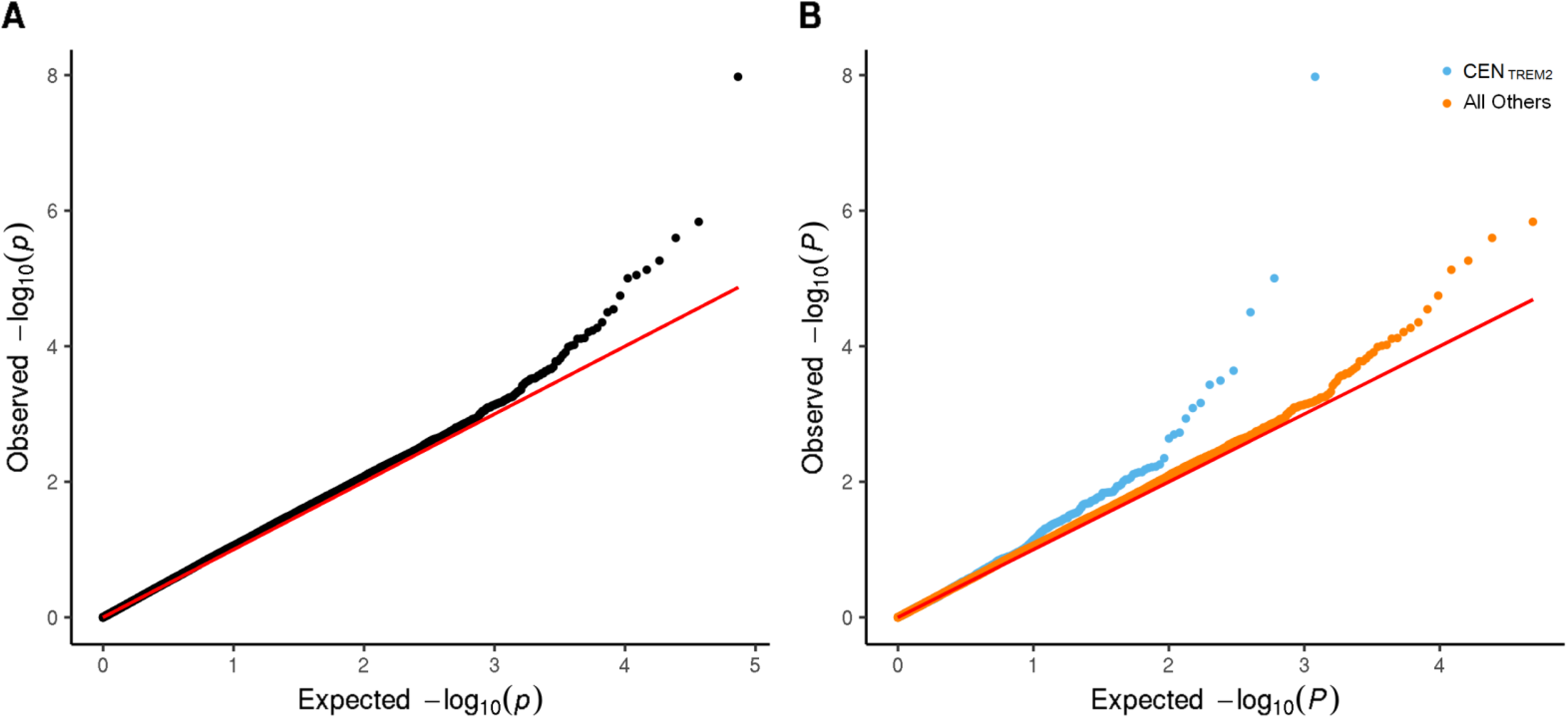
Q-Q plots of P_ADSP_-values for ADSP WES variants with minor allele counts (MAC) of 20 or more. P_ADSP_-values were obtained by logistic regression with appropriate covariates using all 9904 ADSP subjects. Q-Q plots, in which the solid red line shows the P-value distribution expected on the null hypothesis of no association with AD, are shown for P_ADSP_-values of variants with MAF = 0.1% in All Genes of ADSP WES dataset. P_ADSP_-values are shown for all 73,445 variants (MAC ≥ 20) in the 16,310 genes of the ADSP WES dataset. **B** Genes in CEN_TREM2_. P_ADSP_-values are shown for the 1200 variants in the 234 genes of the co-expression network containing *TREM2* (CEN_TREM2_) (blue symbols). For comparison, P_ADSP_-values are shown for the 72,245 variants in the 16,076 genes which remain after CEN_TREM2_ genes are removed from the ADSP WES dataset. These remaining variants (red symbols) also show significant (P_PGS_ = 5.23E-03) association with AD by Broad/WashU PGSA.

### Cumulative Broad/WashU PGSA of CEN_TREM2_ by gene

To evaluate the individual genes in CEN_TREM2_, Broad/WashU PGSA was employed to test the polygenic scores for all variants (MAF > 0.1%) in each gene for association with AD. The 234 CEN_TREM2_ genes were then ranked by their gP_Br/Wa_–values and tested cumulatively for association with AD by Broad/WashU PGSA (Fig. 3 A, B). This analysis identified a highly significant polygenic component (Pcm = 2.08E-09) composed of 36 variants in 5 genes (g5v36) with Pgene–values ≤ 1.07E-02 that improved the AUC by 1.43%.

**Figure 3.**
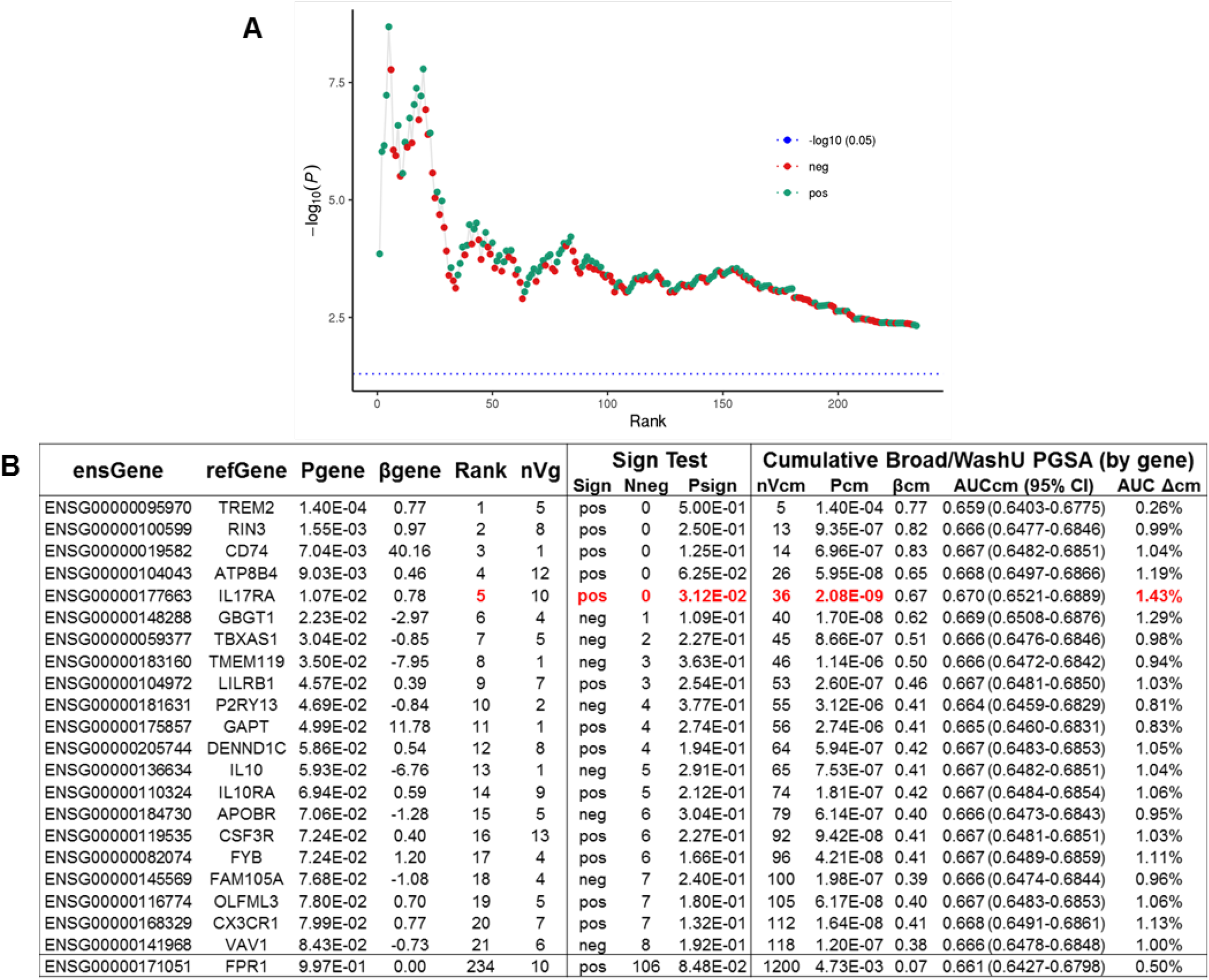
Cumulative PGSA of CEN.TREM2 by gene. All variants in each CEN_TREM2_ gene (nVg) were analyzed by Broad/WashU PGSA to obtain Pgene (gP_Br/Wa_) and βgene (gβ_Br/Wa_). Genes ranked by Pgene were then analyzed by cumulative Broad/WashU PGSA. **A**. Graph shows −log_10_ cumulative P vs Rank. Note that cumulative significance declined whenever the variants in genes with negative gβ_Br/Wa_ (neg, red symbols) were added. B. Tabulated results. Sign test results are for a one sided test analyzing whether the number of positive gβ_Br/Wa_ values (Npos = Rank-Nneg) is significantly greater than the 50% expected on the null hypothesis of no association. Cumulative results show cumulative number of variants (nVcm), as well as cumulative P_PGS_ (Pcm) and β_PGS_ (Bcm). AUC is the AUC and 95% CI for a model including the cumulative polygenic score with *APOE* ε4 dose, *APOE* ε2 dose, and three principal component vectors as covariates. AUC Δcm shows the improvement in AUC in this model compared to a covariates only model, which had an AUC (95%CI) of 0.656 (0.6376-0.6749). Psign and Pcm were most significant and AUC Δcm was maximal when polygenic scores for the 36 variants in the top 5 genes were tested (bold red font).

In addition to P-values for each gene, PGSA generates β-values estimating PGS effect size for each gene (βgene). These βgene-values are directional. If, for example, a gene has a negative value for βgene, then positive polygenic scores derived from Broad βs are associated with *decreased* risk of AD in WashU subjects and vice versa. For such genes, polygenic scores based on Broad βs provide no evidence for association with AD as they do not associate with AD in the predicted direction in WashU subjects. On the null hypothesis of no association, the expectation is that 50% of genes will have positive βgene-values indicating association in the expected direction in test subjects and 50% will have negative βgene-values indicating association in the opposite direction. On the null hypothesis, Pgene-values are distributed uniformly between 0 and 1 so, on average, genes with positive association will be negated by those with negative association resulting in no evidence of association. In CEN_TREM2_, all 5 of the top genes have positive P-values (Fig. 3A, B), and a one-sided sign test shows a significant excess of genes with positive gβ_Br/WaBa_-values over the 50% expected on the null hypothesis (Psign = 3.12E-02).

On the null hypothesis, the 5 most significant genes will sometimes show highly significant association when, by chance, 4 or 5 of the most significant genes have positive βgene-values. Thus our cumulative PGS analysis does not maintain a correct type I error rate when the first five genes are tested. For this reason, results are presented below wherein we evaluate PGS for these genes in the independent Baylor data set aside for follow-up.

### Cumulative PGSA of variants in the 5 gene polygenic component (g5v36)

To evaluate the 36 variants in the 5 gene polygenic component, they were ranked by their Broad P-values (P_Broad_) and tested by cumulative Broad/WashU PGS (Fig. 4A, B). This analysis identified a significant polygenic component (P_PGS_ = 1.43E-09, Psign=9.0E-05) composed of 28 variants with P_Broad_ < 0.56 that improved the AUC by 1.47%. The remaining 8 variants did not show significant association. Thus removal of 8 non-contributing variants with P_Broad_ = 0.56 resulted in a refined polygenic component with 28 variants but P_PGS_ (1.53E-09 vs. 2.09E-09) and the improvement in AUC (1.47% vs. 1.43%) were only slightly better.

**Figure 4.**
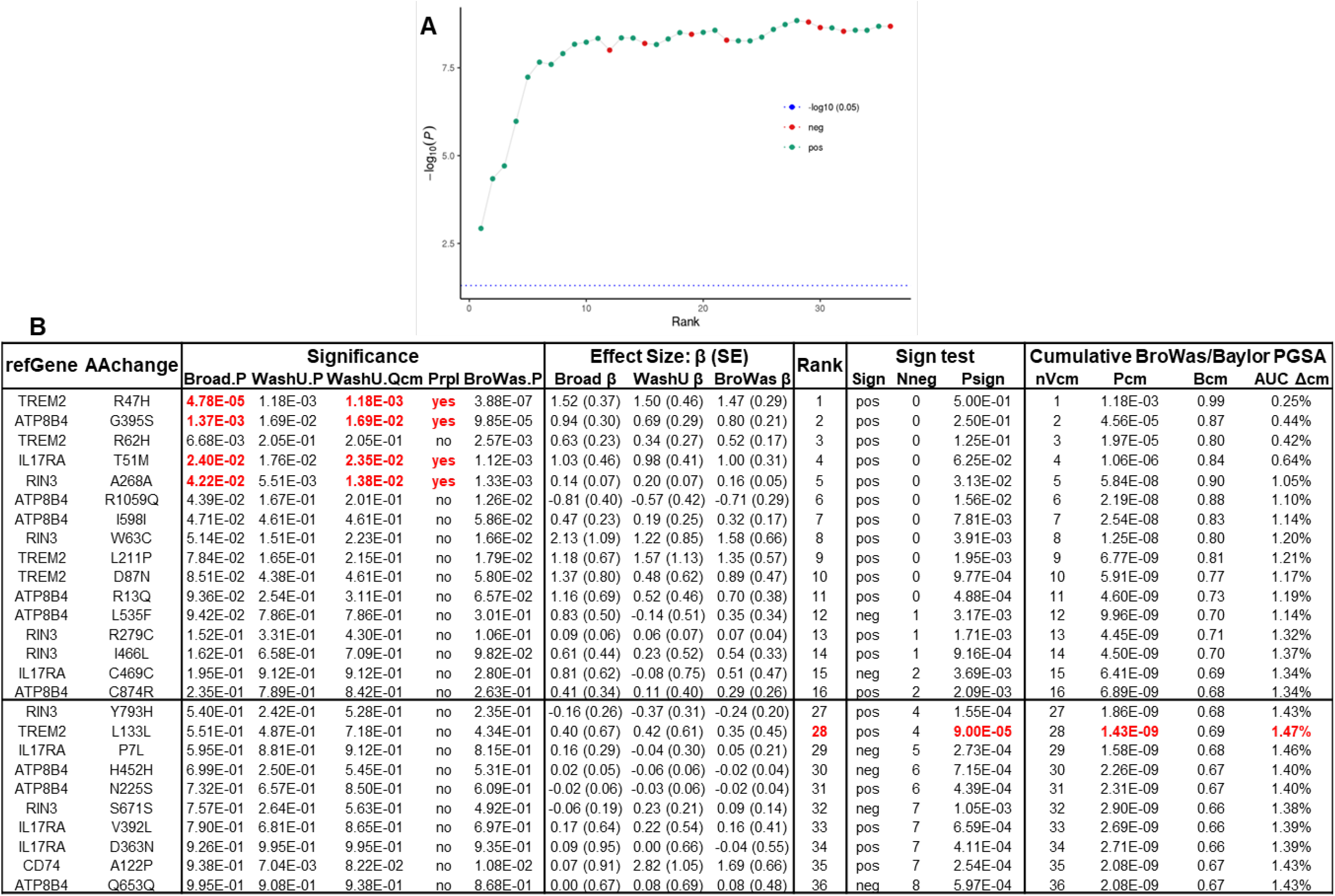
Broad/WashU PGSA of g5v36 variants. **A**. Variants ranked by Broad.P were analyzed by cumulative Broad/WasBay PGSA. A. Graph shows −log_10_ cumulative P vs Rank. Note that cumulative significance declined when (Broad.β * WasBay.β) was negative (neg, red symbols), indicating that the direction of association for the variant added was opposite in Broad and WasBay samples. **B.** Tabulated results. Sign test and cumulative results are tabulated as described in the legend to Fig.3. AUC.Δ was maximal when polygenic scores for the top 28 variants were tested (bold red font). In addition to cumulative PGSA, individual variants, ranked by their P-value in discovery samples (Broad.P), were tested sequentially for significant association in the WasBay samples, adjusting for multiple testing as each variant was tested (WasBay.Qcm). As described in the text, 4 variants showed significant, replicable association with AD (red symbols). Significance (P) and effect size (β) are shown for Broad (discovery) Washu (test) and combined Broad +WashU (BroWas) data. Annotation shows the gene name (refGene: RefSeq gene symbol) and amino acid change (AAchange).

### Variants in g5v36 show significant, replicable association

Of the 36 variants in g5v36, there were 7 variants with nominally significant association in the discovery data (P_Broad_ < 0.05). To identify variants that also showed significant association in the test data, we ordered these variants by their P_Broad_ values and searched sequentially through the 7 variants (Fig. 4B). To adjust for multiple testing, we determined the false discovery rate (FDR)-corrected Q-value (WashU.Qcm) for each variant in the test data as it was evaluated. Searching in this prioritized manner, we found 4 variants that showed significant association in the same direction in both the discovery and test sets (Fig. 4B). The last of these 4 significant variants was found when the variant ranked 5 was tested. Thus, by testing only 5 of the 36 variants in g5v36, we were able to identify 4 variants that showed significant, replicable association with AD (Prpl = yes, Fig. 4B). In the discovery (Broad) data, these 4 variants were the most significant variant in *TREM2* (4.78E-05), *ATP8B4* (1.37E-03), IL17RA (2.40E-02) and RIN3 (4.22E-02).

### PGSA of PCg23 subsets

. The 4 variants that showed significant, replicable association formed a polygenic component (g4v4) that showed highly significant association by Broad/WashU PGSA (P_PGS_ = 1.10E-07) and improved the AUC by 1.15%. The polygenic component formed by the remaining 31 variants in these 4 genes (g4v31) was also significant (P_PGS_ = 1.01E-03), providing independent evidence that these genes associate with AD. Moreover all 4 genes had significant gP_Br/Wa_ values ranging from 5.40E-06 to 3.19E-02. The polygenic component formed by all 35 variants in the 4 genes (g4v35) was highly significant (P_PGS_ = 2.72E-09) and improved the AUC by 1.39%. Among the 4 variants that showed significant, replicable association, the most significant is the well-established *TREM2* p.R47H variant. Another variant is in *RIN3*, which has previously been linked to AD as it is in a region tagged by a significant GWAS SNP (rs10498633)^20^ that also includes *SLC24A4*. The remaining variants are in novel genes not previously linked to AD (*ATP8B4*, *IL17RA*).

### Follow-up analysis of g4v35 by BroWas/Baylor PGSA

To test g4v35 variants in independent case-control samples, polygenic scores were analyzed in the Baylor data set aside for follow-up. To optimize this analysis, variants were analyzed by logistic regression using combined Broad and WashU data, and BroWas-derived polygenic scores were tested for association with AD in the Baylor data. By BroWas/Baylor PGSA (Fig. 5A), polygenic scores for all variants in g4v35 showed significant association (P_PGS_ = 4.31E-04) and improved the AUC by 0.43%

**Figure 5.**
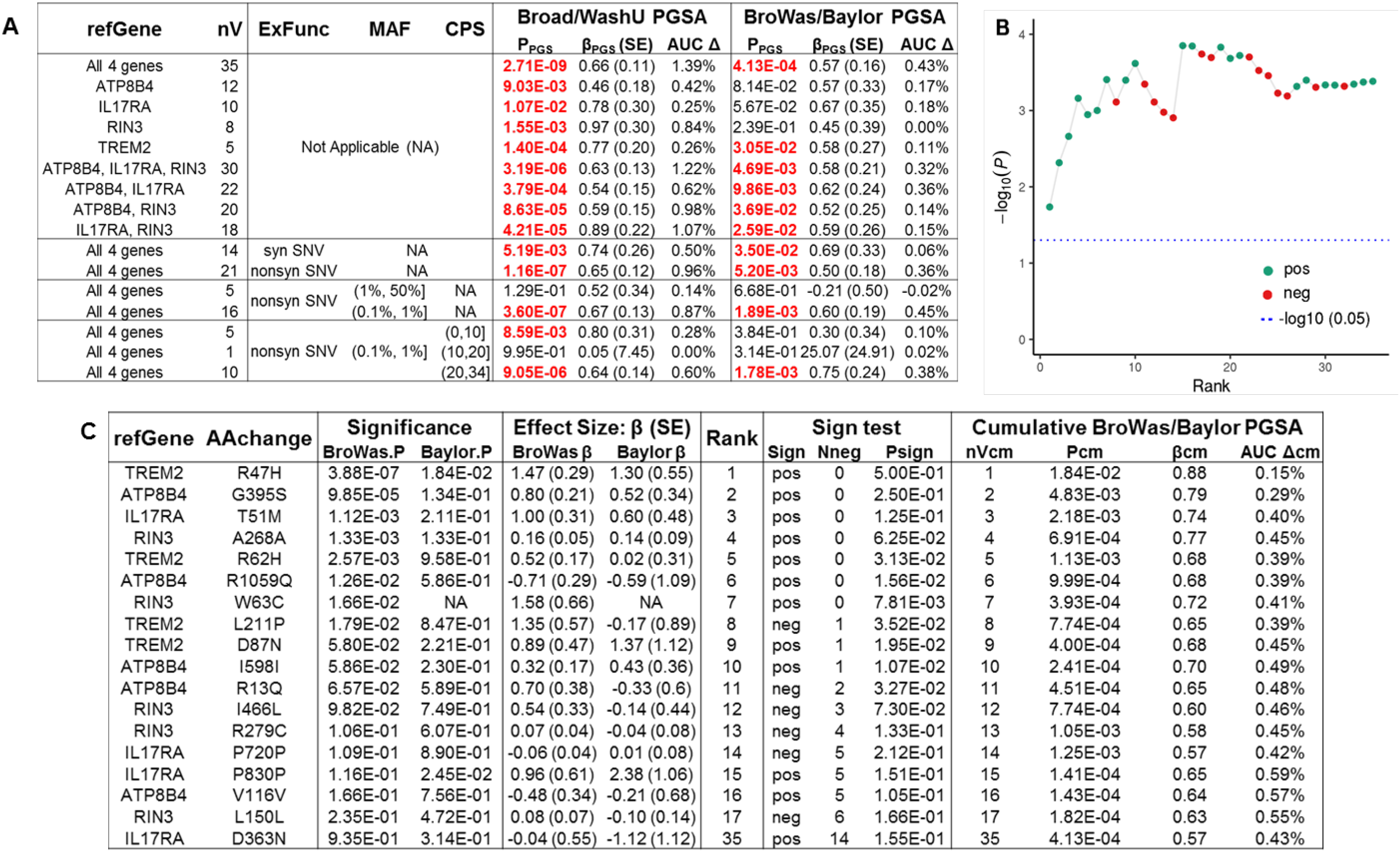
BroWas/Baylor PGSA of g4v35 variants. **A.** BroWas/Baylor PGSA of g4v35 compared with Broad/WashU PGSA. To test g4v35 variants in independent case-control samples, polygenic scores were analyzed in the Baylor data set aside for follow-up. To optimize PGSA, variants were analyzed by logistic regression using combined Broad and WashU data, and BroWas-derived polygenic scores were tested for association with AD in the Baylor data. Comparative results are shown for all 4 genes, each gene, and several combinations of genes. Comparative results are also shown after stratification by exonic function (ExFunc) which compared synonymous SNV (syn SNV) with non-synonymous SNV (nonsyn SNV), minor allele frequency (MAF), and CADD PHRED-scaled score (CPS). **B.** Variants ranked by BroWas.P were analyzed by cumulative BroWas/Baylor PGSA. Graph shows −log_10_ cumulative P vs Rank. Note that cumulative significance declined when (BroWas.β * Baylor.β) was negative (neg, red symbols), indicating that the direction of association for the variant added was opposite in BroWas and Baylor samples. **C.** Tabulated results. Sign test and cumulative results are tabulated as described in the legend to Fig.3. AUC.Δcm was maximal and Pcm was most significant when polygenic scores for the top 15 variants were tested. Annotation shows the gene name (refGene: RefSeq gene symbol) and amino acid change (AAchange) if any.

When we considered the gene-level PGS for each gene in g4v35 (Fig. 5A), only the PGS for *TREM2* showed significant association with AD (P_PGS_ = 0.031). The gene-level PGS for the other 3 genes had βgene-values in the expected direction but were not significant (*ATP8B4* P_PGS_ = 0.081; *IL17RA* P_PGS_ = 0.051; and *RIN3* P_PGS_ = 0.24). However, the PGS constructed using the 30 variants from these genes showed significant association (P_PGS_ = 4.69E-03) with AD. Furthermore, PGS for the *ATP8B4*-*IL17RA* pair (P_PGS_ = 9.88E-03), ATP8-RIN3 pair (P_PGS_ = 3.69E-02), and IL17RA-RIN3 pair (P_PGS_ = 2.59E-02) were also significant (Fig. 5A).

The 35 variants in g4v35 were also analyzed in the Baylor data (Fig. 5B) by ranking them according to their P_BrWa_-values and performing cumulative BroWas/Baylor PGSA. This analysis identified a polygenic component (g4v15) composed of the 15 most significant variants that showed significant association with AD (P_PGS_ = 1.40E-04) and improved the AUC by 0.59%.

### PGSA of g4v35 stratified by exonic function, MAF and deleteriousness

To evaluate the variants in g4v35 further, they were analyzed by Broad/WashU PGSA and follow-up BroWas/Baylor PGSA after stratification by exonic function (Fig. 5A). In g4v35, association was driven primarily by the 21 non-synonymous SNVs. Polygenic scores for these non-synonymous variants were significant both by Broad/WashU PGSA (P_PGS_ = 1.16E-07) and on BroWas/Baylor follow-up (P_PGS_ = 5.20E-03). Although less significant, the 14 synonymous SNVs were also significant by Broad/WashU PGSA (P_PGS_ = 5.19E-03) and on BroWas/Baylor follow-up (P_PGS_ = 3.50E-02).

Further stratification of the 21 non-synonymous SNVs by MAF showed that association was driven by low frequency variants with MAF of 0.1 to 1%. The 16 low frequency, non-synonymous variants were significant both by Broad/WashU PGSA (P_PGS_ = 3.60E-07) and on BroWas/Baylor follow-up (P_PGS_ = 3.50E-02), whereas the 5 higher frequency variants with MAF of 1 to 50% were not significant by Broad/WashU PGSA (P_PGS_ = 1.29E-01) or BroWas/Baylor follow-up (P_PGS_ = 6.68E-01).

Analysis of the 16 low frequency non-synonymous SNVs after stratification by their CADD^21^ PHRED-scaled scores (CPS) showed that association was driven primarily by the 10 variants with CPS of more than 20 estimated to be highly deleterious (Fig. 5A). These variants showed significant association with AD both by Broad/WashU PGSA (P_PGS_ = 9.05E-06) and on BroWas/Baylor follow-up (P_PGS_ = 1.78E-03). The 5 variants with CPS of 10 or less estimated to be less deleterious were significant by Broad/WashU PGSA (P_PGS_ = 8.59E-03) but not by BroWas/Baylor follow-up (P_PGS_ = 3.84E-01).

### Sequence kernel association testing (SKAT-O)

Analyses using SKAT-O^22,23^ provide an additional opportunity to test the 4 genes in g4v35 for association with AD because rare variants (MAF ≤ 0.1%) that cannot be analyzed by PGSA can be analyzed by SKAT-O. The 4 genes in g4v35 had 215 variants with MAF ≤ 0.1% (g4v215), and these variants showed significant association by SKAT-O (P_SK_ = 2.87E-03)

### SKAT-O of g7v325 variants stratified by exonic function and deleteriousness

Among the 215 rare (MAF ≤ 0.1%) in g4v215, the 172 variants that alter protein (Fig. 6A) showed significant association with AD (P_SK_ = 3.545E-04) and were composed of 3 stop-gain variants (P_SK_ = 8.77E-02), 153 nonsynonymous SNVs (P_SK_ = 7.36-04) and 16 indels (frameshift and non-frameshift insertions and deletions) (P_SK_ = 3.38E-01). The 43 synonymous SNVs (P_SK_ = 8.58E-01) showed no evidence of association.

**Fig 6.**
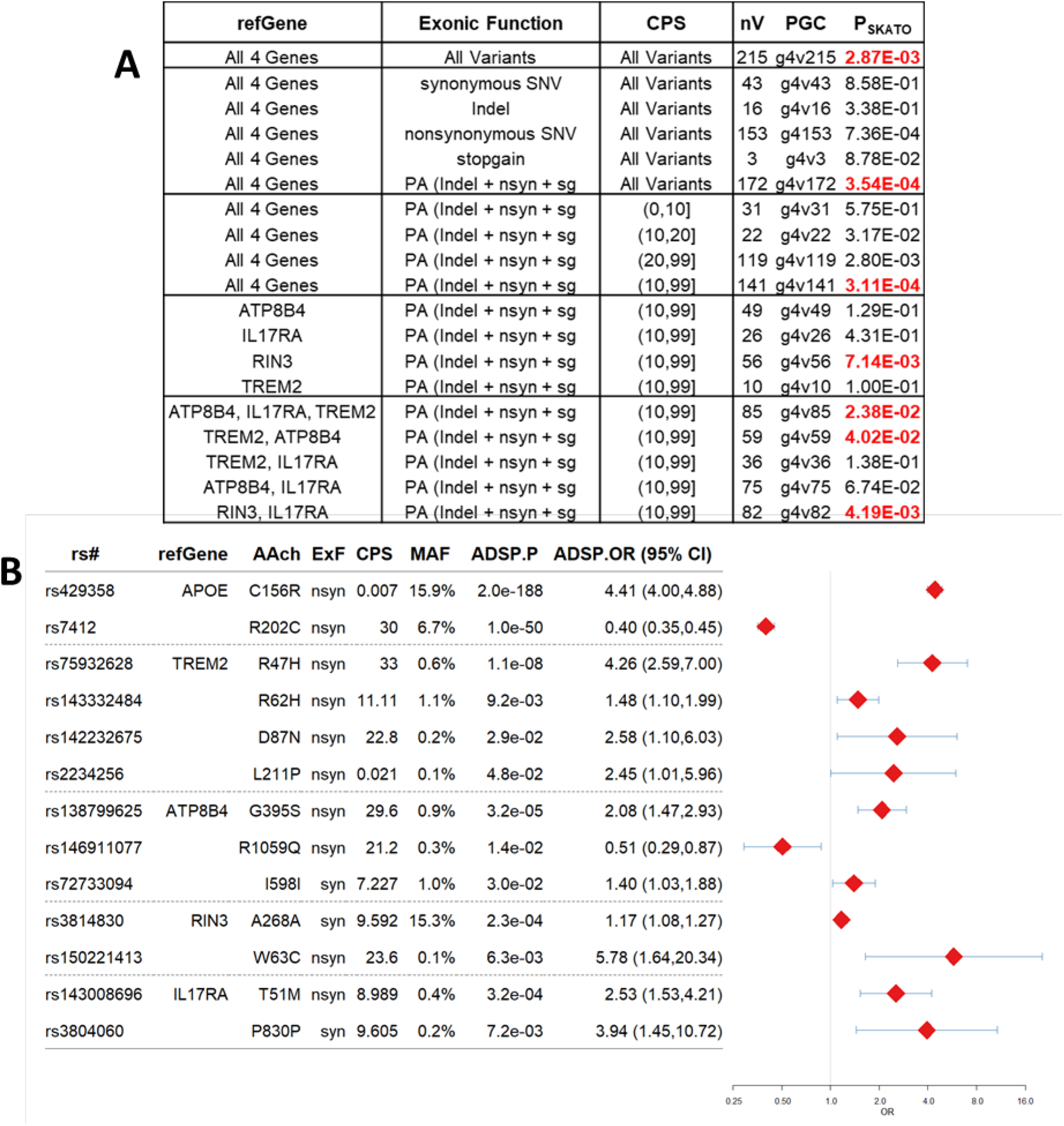
SKAT-0 analysis of g4v215and Forest Plot of g4v11. **A.** SKAT-O results for polygenic components composed of variants with MAF ≤ 0.1%. Results are shown for all 215 variants with MAF ≤ 0.1% in the 4 four genes identified by PGSA, for stratified analysis of g4v215 by exonic function and CADD PHRED-scaled score, for individual genes, and for gene combinations. **B.** Forest plot of variants in *TREM2*, *ATP8B4*, *RIN3* and *IL17RA* with ADSP.P-values < 0.05. For comparison results are shown for the APOE SNPs that tag the APOE ε4 and .ε4 alleles. Annotation: rs# (dbSNP_id_142), AAch (Amino acid change) and ExF (exonic function per Ensembl gene), CPS (CADD PHRED-scaled score). Symbol size is proportional to the number of samples in which genotypes were determined (9904 ADSP samples for all variants)

Analysis of the 172 protein-altering (PA) variants (Fig. 6A) after stratification by their CADD^21^ PHRED-scaled scores (CPS) showed that association was driven primarily by the 119 variants with CPS of more than 20 estimated to be highly deleterious (P_SK_ = 2.80E-03) and the 22 variants with CPS of 10-20 estimated to moderately deleterious (P_SK_ = 3.17E-02). The 31 variants with CPS of 10 or less showed no evidence of association (P_SK_ = 5.75E-01). Thus the association of g4v215 variants with AD was due to 141 protein altering variants (g7v141) with CPS > 10 (P_SK_ = 3.11E-04).

Analysis of these 141 variants by gene showed that the 56 variants in RIN 3 (P_SK_ = 7.14E-03) were significant. When the 10 variants in *TREM2* (P_SK_ = 1.00E-01) and the 49 variants in *ATP8B4* (P_SK_ = 1.28E-01) were tested together, the combined set of 59 variants in the two genes was significant by SKAT-O (P_SK_ = 4.02E-02). The 26 variants in IL17RA (P_SK_ = 4.31E-01) were not significant, but the 85 variants in the combined set of TREM2, ATP8B4, and IL17RA showed improved significance (Psk=2.37E-02) compared to TREM2 and ATP8B4, the combined set of 82 variants in IL17RA and RIN3 (Psk=4.18E-03) showed improved significance compared to RIN3 alone, and the 75 variants in IL17RA and ATP8B4 ((Psk=6.74E-02) showed improved association that was more significant than *ATP8B*4 alone.

## Discussion

In this study we begin by using Broad/WashU PGSA to show that polygenic scores for all pruned ADSP variants and for the 1200 variants in CEN_TREM2_ are significantly associated with AD (Fig.1). We then test the hypothesis that among the 234 genes in CEN_TREM2_ there will be some with a variant that shows significant association at α = 0.05 in both the discovery (Broad) and test (WashU) data. Genes are a logical way to organize WES data by function, and it is well established that genes with variants that cause or alter risk of disease typically have many such variants. Thus, for example, the *APP*, *PSEN1*, and *PSEN2* genes all have multiple variants that cause early onset familial AD, and *APOE* has two powerful variants which form three haplotypes that alter risk of AD. We reasoned, therefore, that analysis by gene might capture exonic variants associated with AD better than analysis by variant. More specifically, this reasoning suggested that genes with a significant, replicating variant were likely to be found among the genes with polygenic scores that associated most significantly with AD.

For this reason we began our analysis of CEN_TREM2_ by ordering genes by their gP_Br/Wa_–values and performing cumulative Broad/WashU PGSA (Fig. 1, Fig. 3). This analysis showed that association became most significant when the 5^th^ gene was tested. This result was significant by one-sided sign testing because the top 5 genes all had positive gβ_Br/Wa_-values (Psign = 3.25E-02). Empirical testing showed that polygenic scores for the top 5 genes also showed significant association with AD (empP_PGS_ < 2.6E-03). The top 5 genes had only 36 variants, among which 7 (19%) had P_Broad_-values <= 0.05. These variants were ranked by their P_Broad_ values and tested sequentially for significant association in the test (WashU) data, adjusting for multiple testing by determining the false discovery rate (WashU.Qcm) in the test data as each variant was tested. For 4 genes (*TREM2*, *ATP8B4*, *RIN 3*, *IL17RA*), the most significant variant in the discovery data had a P_Broad_-value < 0.05 and a WashU.Qcm-value < 0.05 (Fig. 1, Fig. 4). By itself, this result provides strong evidence that the most significant variant in each gene associates with AD. Polygenic scores for the remaining 31 variants (g4v31) also showed significant association with AD, providing additional evidence that these 4 genes associate with AD.

The 5^th^ gene (*CD74*) in the set of 5 that were most significant is informative. This gene had only 1 variant with a MAF > 0.1%. Both the gene (Fig. 3B) and variant (Fig. 4B) were significant in the WashU data (P_WashU_ and gP_Br/Wa_ = 7.04E-03), but in the Broad data (Fig. 4B) this variant showed no evidence of association with AD (P_Broad_ = 9.38E-01). Thus, the highly significant association of the CD74 variant with AD in the WashU data is likely occurring primarily, if not exclusively, by chance alone. In sharp contrast to the other variants, which all showed significant association in both the WashU and Broad data, this was evident when association of the CD74 variant was examined in the Broad data where there was no evidence of association with AD.

The sequential analyses (Fig. 1) performed using Broad samples for discovery and WashU samples for testing provide compelling evidence that g4v35, a polygenic component composed of exonic variants in *TREM2*, *ATP8B4*, *RIN3*, and *IL17RA*, shows significant association with AD wherein the most significant variant in each gene shows powerful association that is significant in both the discovery and test data. These results establish that there are AD genes with powerful variants within g4v35, but they do not establish that each gene in the polygenic component is an AD gene. However likely or unlikely it may seem to someone reviewing the results for each of these genes, there is always the possibility that a gene within g4v35 may be associating with AD by chance alone in the Broad and WashU data. At the beginning of this analysis, we identified 16,308 genes in the ADSP dataset. With that many genes, there are bound to be some genes with variants that, by chance alone, show association with AD that closely resembles the association observed in a true AD gene. For this reason, and because the approach used to identify g4v35 was unconventional, it was important to perform follow-up analyses to confirm that these 4 genes associate with AD. We did this by analyzing independent Baylor subjects by BroWas/Baylor PGSA, and by analyzing independent variants with MAF ≤ 0.1% by SKAT-O.

BroWas/Baylor PGSA (Fig. 1, Fig. 5A) confirmed that polygenic scores for the 35 variants in the four genes (g4v35) show significant association with AD (P_PGS_ = 4.13E-04) and that association is driven primarily by the 21 non-synonymous variants forming g4v21 (P_PGS_ = 5.20E-03) with a significant contribution from the 14 synonymous variants comprising g4v14 (P_PGS_ = 3.50E-02). BroWas/Baylor PGSA also confirmed that the association of non-synonymous variants was driven primarily by the 10 deleterious (CPS > 20), non-synonymous variants with MAF of 0.1 to 1.0% comprising g4v10 (P_PGS_ = 1.78E-03).

Because there are fewer samples in the Baylor data than in the WashU data, our expectation was that polygenic scores for each gene would show less significant association in the Baylor data than in the WashU data, where all 4 genes were significant. This did, in fact, occur (Fig. 5A). *TREM2* (gP_BrWa/Ba_ = 3.05E-02) continued to be significant. Though not significant at α = 0.05, *IL17RA* (gP_BrWa/Ba_ = 5.67-02), *ATP8B4* (gP_BrWa/Ba_ = 8.14-02), and RIN3 (gP_BrWa/Ba_ = 2.39E-01) showed suggestive association with AD. That each of these 3 genes associated with AD on follow-up analysis of Baylor samples was evident when polygenic scores for the 3 pairs of genes [(ATP8B4, IL17RA); (ATP8B4, RIN3); (IL17RA, RIN3)] formed by these genes were analyzed. By BroWas/Baylor PGSA, polygenic scores for each pair showed significant association with AD even though polygenic scores for each gene showed only suggestive association (Fig. 5A).

By SKAT-O, the 141 deleterious, protein-altering variants with MAF ≤ 0.1% in the four genes (g4v141) showed significant association with AD (P_SK_ = 3.11E-04) in the 9904 samples comprising the ADSP data (Fig. 6A). Of these 141 variants, there were 53 in *RIN3* that showed significant association with AD (P_SK_ = 7.14E-03). The 10 variants in *TREM2* (P_SK_ = 1.00E-02) and the 49 in *ATP8B4* (P_SK_ = 1.29E-01) showed suggestive association with AD that became significant (P_SK_ = 4.02E-02) when the combined 59 variants were analyzed (Fig. 6A). Thus *TREM2* and *ATP8B4* have deleterious, protein-altering variants with MAF ≤ 0.1% that contribute to significant association with AD. The 26 variants in IL17RA (P_SK_ = 4.31E-01) were not significant, but when these variants were added to the 59 variants in *TREM2* and *ATP8B4*, the combined set of 85 variants showed improved significance (P_SK_ = 2.38E-02 vs 4.02E-02) suggesting that deleterious, protein-altering variants with MAF ≤ 0.1% in *IL17RA* may also show weak, non-significant association with AD.

Among the 35 variants in the 4 genes, 11 (31%) showed nominally significant association with AD (P_ADSP_ < 0.05), and 10 of these were low frequency variants (MAF 0.1 to 1.0%) associated with strongly increased (9) or decreased (1) risk of AD comparable to that of the well-known *APOE* ε4 and ε2 alleles. This is illustrated in the well-annotated Forest plot of Fig. 6B, which shows the OR and 95% CI for these 11 variants in ADSP samples. For reference, the SNPs tagging the *APOE* ε4 and ε2 alleles are shown at the top of Fig. 6B. There were multiple variants with ADSP.P-values < 0.05 in *TREM2* (4) and *RIN3* (2), two genes previously associated with AD and in *ATP8B4* (3) and *IL17RA* (2). Neither *ATP8B4* nor *IL17RA* were linked to AD by the AD GWAS performed to date^20,24,25^, but Holstege, et al^26^ recently reported that carrying rare damaging variants in *ATP8B4* is associated with AD.

Like *TREM2*, *ATP8B*4 and *IL17RA* are selectively expressed in human microglial cells^7^ (Supplementary Fig. 2). Like the 4 nominally significant variants in *TREM2*, the 5 nominally significant variants in *ATP8B4* and *IL17RA* have MAF of 0.1 - 1.0%. The 5 variants in these genes are associated with strongly increased (4) or decreased (1) risk of AD. Genes like these are ideally suited for studies aimed at understanding how exonic variants modulate microglial function to increase or decrease risk of AD. Importantly, the effect(s) of these variants may occur in the window of immunomodulatory opportunity wherein Aβ oligomerization and deposition have occurred, are detectable, and have prompted a microglial response but cognitive decline has not yet begun.

## Methods

### ADSP WES Variant Calling

Samples in the ADSP data set^19^ were sequenced at three large scale sequencing and analysis centers (LSACs) located at the Broad Institute (Boston, MA), Baylor College of Medicine, (Houston, TX) and Washington University (St. Louis, MO). The Baylor and Washington University LSACs used the Nimblegen VCRome.2.1 exome capture kit (35.3Mbp); the Broad Institute used Illumina’s Rapid Capture kit (37.7Mbp). Following IRB approval and DUC agreement, whole exome sequencing (WES) data and related phenotypes for 10933 samples from the ADSP WES case-control study spanning 6 cohorts (phs000572.v4.p2) were downloaded from dbGaP. Four samples that were either not part of or retracted from the ADSP in subsequent data releases were removed from our analyses. The most up-to-date phenotypes and sample information (phs000572.v7.p4) were used to analyze the remaining 10,929 samples.

WES files (fastq) obtained from dbGaP were processed with GenomeGPS (v3.0.1), a comprehensive secondary analysis pipeline for sequencing data at Mayo Clinic. Reads were aligned to the reference genome (hg19) using Novoalign^18^ (args: *-x 5 -g 40 -i PE 425,80 -r Random --hdrhd off -v 120*). Quality of sequencing reads was assessed using FastQC^27^. After marking duplicates using Picard^28^ tools, variant discovery and genotyping were carried out with genome analysis toolkit ^29^ (GATK) v3.3 and implemented using GATK’s Best Practices workflow. After realignment and recalibration, variant calling on each sample (SNPs and INDELs, simultaneously) was performed using GATK’s *HaplotypeCaller*. Joint genotyping of variants in common capture regions (regions common to both capture kits used by the three LSACs ~ 34Mbp, identified using bedtools^30^) across all samples was performed using GATK’s *GenotypeGVCFs* to generate a consensus variant call file. Quality of SNPs and INDELs was assessed separately using GATK’s *VariantRecalibrator* (**SNP**: “*-an QD -an MQRankSum -an ReadPosRankSum -an FS*”, **INDEL**: “*-an QD -an FS -an ReadPosRankSum --maxGaussians 4*”) and *ApplyRecalibration* (ts_filter: 99.0) tools, a process known as variant quality score recalibration (VQSR).

### Sample Quality Control

#### Read coverage

Read coverage of the exome capture region was assessed for each sample. A commonly-used threshold for high-quality exome sequencing with sufficient depth would exclude any sample with less than 50% of the capture region covered at 40X. This threshold would remove a group of otherwise high-quality samples, so we lowered the threshold to retain samples with high coverage at 10X, yet lower coverage at 40X. Samples with less than 90% of the capture region covered at 10X, or less than 30% covered at 40X were excluded. Samples with less than 50% coverage at 40X were flagged and investigated for other QC metrics. Similarly samples with a call rate of less than 95% for SNVs or 90% for INDELs or missing chromosomes were flagged.

#### Sex

To verify the sex of samples, clinical information provided by the ADSP was compared to genotypes on the sex chromosomes. Variants on the X chromosome that passed VQSR, had a minor allele frequency greater than 0.002, call rate of 95% and above and a Hardy Weinberg p-value greater than 1e-08 were used to assess sex. Variants in the pseudo autosomal regions of the X-chromosome were excluded from the analysis. Using PLINK v1.9^31^, variants were pruned to an r-squared of 0.05 within a sliding window of 50 variants (--indep-pairwise 50 5 0.05). The resultant homozygosity estimate (F) of the X-chromosome for males and females was used to exclude samples. A measure closer to 1 is expected for males and closer to 0 for females. Samples marked females having an F estimate of 0.3 or less and males with an F-estimate equal to or greater 0.7 were retained. For those with an F estimate opposite to what was expected, it is likely that sex was mis-classified in the clinical variables, or that there was a sample mix-up. Samples closer to the thresholds were examined for other QC issues.

#### Ti/Tv ratio

The transition to transversion (Ti/Tv) ratio was examined for each sample using all variants in the common capture region and also for the subset of common capture, exonic SNPs that passed VQSR. For coding variants, Ti/Tv ratios are expected to be around 2.8. The distribution of Ti/Tv ratios for all variants in common-capture regions centered around 2.5, but the common capture, exonic SNVs that passed VQSR had a Ti/Tv ratio of greater than 2.75 and only 27 samples had a Ti/Tv ratio of less than 2.8. Hence no samples were excluded under this metric.

#### Sample contamination

Contamination between samples was examined using *VerifyBamID*^32^, a tool that checks whether reads in sample match previously observed genotypes in another sample (or a group of samples). We applied the sequence-only method of VerifyBamID, which estimates contamination by modeling the sequence reads as a mixture of two unknown samples based on the allele frequency information provided in a reference VCF file. The 1000genomes array genotypes were used as reference for this analysis. Given the sample size, contamination estimation was executed as a two-step process. Initially VerifyBamID was run on chromosome 20 for all samples. A FREEMIX score (a VerifyBamID sequence-only contamination estimate), of 0.02 was used as a threshold to identify samples with potential contamination. For those samples with suspected contamination, VerifyBamID was run on all chromosomes (whole exome). Samples with a whole exome FREEMIX score greater than 0.04 were excluded. Samples with a FREEMIX greater than 0.02 for chromosome 20 but less than 0.04 for all chromosomes, showing some level of contamination, were examined for other QC issues. A large portion of samples that failed sex-check were removed for contamination as well.

After evaluating sample quality using the metrics mentioned above, a total of 10715 samples passed QC. 25 samples were excluded for insufficient read coverage (19 failed for having < 30% of the capture region covered at 40x and 6 samples failed for having <90% covered at 10x). 29 samples failed to meet the call rate threshold of 95%. 26 samples were identified as having missing chromosomes and 143 samples were excluded for having a whole genome FREEMIX score 0.04 or greater, showing significant levels of sample contamination. Of these, 133 were sequenced at Baylor, 1 at the Broad and 9 at Washington University. 68 samples with a homozygosity estimate for females greater 0.3 and males less than 0.7 were also excluded. Some of the samples that were excluded failed in more than one metric.

#### Relatedness

Relatedness among samples in the ADSP cohort was examined using KING-robust^33^, a tool to identify relationships by estimating pairwise kinship coefficients and identity by state probabilities using genotype data. KING is able to clearly delineate unrelated samples from those that are related, up to the 3^rd^ degree, and is robust to population substructure. Only samples that passed aforementioned QC measures were used to estimate kinship coefficients. These kinship coefficients along with an IBS0 score (the probability of sharing 0 variants identical by state, which is lower for closely related pairs), were used to identify related samples. Initially 29 samples with a kinship coefficient greater than 0.3 and an IBS0 close to 0 were identified. Of these, 9 samples were identified to have been sequenced in triplicates and one sequenced in duplicate. From these 29 samples, 10 high quality samples were retained and 19 were removed. Thus, after removing 19 from the set of 10715 samples identified above, the remaining 10696 samples were reprocessed for multi-sample calling and joint genotyping.

After joint genotyping and VQSR of the 10696 high quality samples, analysis with KING-robust was repeated to identify and remove additional related samples. As a first step, groups of related samples with a kinship coefficient equal to or greater than 0.0442 were identified. Subjects in these “families” were prioritized to keep the least contaminated sample obtained from patients with AD. If any of these samples in a family were grouped together as a result of underlying contamination, all samples in the group were excluded. For every pair of related samples with a kinship coefficient greater than or equal to 0.0442 (1^st^, 2^nd^ and 3^rd^ degree relatives), the sample with lower levels of contamination (FREEMIX for whole genome less than 0.02), obtained from a subject with AD and having better coverage metrics (in that order of precedence), was chosen to be retained. In summary, 42 samples were dropped at the 1^st^ degree (kinship coefficient: 0.177-0.354), 9 samples were dropped at 2^nd^ degree (0.0884-0.177) and 76 samples were dropped at 3^rd^ degree (0.0442-0.0884) of relatedness. A total of 10569 samples were retained after QC for relationship status.

#### Population stratification

In order to retain relatively homogeneous, Caucasian samples of European descent, sample population was evaluated using principal component analysis (PCA). A set of unrelated samples that passed prior QC metrics (n=10569) were examined for population stratification using Eigenstrat^34^. Prior to performing PCA, variants were pre-selected for the following: autosomal SNPs that pass VQSR with a genotyping rate 95% or more, having minor allele frequency greater than 0.01 and meeting a Hardy-Weinberg threshold of 1e-05. In addition, any SNPs associating with any of the LSACs with a p-value greater than 1e-07 were excluded. Variants in highly variable and duplicitous regions of the human genome, along with those in and around ApoE locus were also excluded. Remaining variants pruned to an r^2^ less than 0.1 (nSNPs=15,438) were subsequently used with Eigenstrat to perform PCA. Eigenstrat was set to removes outliers up to 6 standard deviations for the top 10 principal components (PCs) over 6 iterations, while refitting PCs after each iteration of outlier removal. Of 10569 high quality unrelated samples, 10241 were retained.

#### APOE dosage

As an additional QC metric relevant to Alzheimer’s disease, clinical *APOE* genotypes provided with the ADSP samples were compared with the genotypes obtained by WES, and 337 samples with discordant genotypes were eliminated leaving 9904 samples in the final dataset.

### Variant Quality Control

Variants in autosomes passing VQSR FILTER, originating from non-multi-allelic loci and having a genotyping rate of over 95% across all samples were retained. Variants in regions known to lead to spurious associations were excluded. Variants that had a Bonferroni adjusted Hardy Weinberg *p-value* less than 0.05 in controls were also excluded. Additionally, for logistic regression analysis with covariates, variants with minor allele counts of less than 20 were excluded. Due to the known strong association of variants in the APOE locus with AD, variants in the APOE LD block (chr19: 45,000,000-45,800,000bp) were excluded from the polygenic score analysis.

### Variant annotation and PLINK genotypes

Variants were annotated with information from public databases using Annovar^35^. Using PLINK 1.9, the VCF file with variant genotypes was converted to files suitable for subsequent analyses with PLINK 1.9 software. A PLINK compatible covariate file was generated. While converting genotypes from VCF to PLINK, SNPs and INDELs were processed separately. At multi-allelic sites, we let PLINK retain the most common alternate allele. All non-variant sites were dropped. SNP IDs were encoded using chromosome (CHR), position (POS), minor (A1) and major (A2) alleles as “CHR:POS:A1:A2”.

### Single variant analysis, additional QC for LSAC heterogeneity

Single variant analysis was performed on 102,826 exonic variants which passed QC and had a minor allele count (MAC) of twenty or more. Variants were analyzed by logistic regression using an additive model with sex*, APOE* ε4 dose, *APOE* ε2 dose, LSACs, and three principal component vectors as covariates. As a final QC measure, multinomial regression was performed to assess heterogeneity in minor allele frequency across the three LSACs, and 677 variants (0.66%) with study-wide P_LSAC_ values ≤ 4.86E-07 were removed.

### Weighted gene co-expression network analysis

Weighted gene co-expression network analysis (WGCNA) was performed using R package WGCNA^36^ to identify co-expressed genes in an RNA sequencing (RNAseq) dataset. The cohort, generation of RNAseq data and quality control steps have been described previously^18,37^. Briefly, RNA was isolated from temporal cortex tissue of neuropathologically diagnosed AD patients and controls. RNA libraries were generated using the TruSeq RNA Sample Prep Kit (Illumina, San Diego, CA) and sequenced on an Illumina HiSeq2000 (101bp PE) multiplexing 3 samples per lane. Raw reads were aligned to GRCh37 and were counted for each gene through the MAP-RSeq pipeline^38^. Gene read counts were normalized using conditional quantile normalization^39^. After QC, 80 AD and 76 control samples were retained for analysis. To account for covariates, expression residuals were obtained using multiple linear regression implemented in R, where gene expression was the dependent variable, and sex, age at death, flow cell and RNA integrity number (RIN) were the independent variables. As previously described, co-expression analysis was performed for 13,273 TCX RNAseq transcripts (13,211 unique genes), which were expressed above background levels in both this RNAseq dataset and in an independent cohort^37^. Co-expression networks based on residuals were obtained using WGCNA function *blockwiseConsensusModules* (args: networkType=“signed”, TOMType=“signed”, power=12). For genes in each co-expression network, enriched gene ontology (GO) terms were identified by function *GOenrichmentAnalysis*. Eigengenes that represent each co-expression network were obtained from function *moduleEigengenes*. One module was identified to contain *TREM2* and thus genes in this module (CEN_TREM2_) were selected for further study. The CEN_TREM2_ module contained 295 genes, of which 234 had variants in the ADSP dataset that passed QC and were subsequently carried forward for analysis.

### Polygenic score analysis (PGSA)

#### Broad/WashU PGSA of pruned variants (MAC≥ 20) in the ADSP and CEN_TREM2_

To evaluate variants with a MAC ≥ 20 in the entire ADSP dataset, the 234 genes of CEN_trem2_, and the ADSP variants remaining after removing variants in 234 genes of CEN_trem2_, we performed polygenic score analysis (PGSA) with the Broad genotypes as the discovery set and the Washington University genotypes as the test set. Baylor genotypes were reserved for follow-up analysis. The single variants in the discovery set (Broad) were analyzed by logistic regression using an additive genetic model adjusting for sex, *APOE* ε4 dose, *APOE* ε2 dose, and three principal components addressing population substructure. Using the clump function in PLINK v1.9^31^, variants were pruned to reduce linkage disequilibrium (r^2^ < 0.2). The pruned betas estimated by logistic regression were then used to construct polygenic scores for each subject in the test set (WashU) which were tested for association with AD.

#### Gene-level Broad/WashU PGSA of CEN_TREM2_

To evaluate association of each CEN_TREM2_ gene with AD, PGSA was employed to analyze all pruned variants with MAC ≥ 20 in each gene. Broad/WashU PGSA, performed as described above, was used to obtain P_GENE_-values for each gene in samples genotyped at WashU. The 234 CEN_trem2_ genes were then ranked by their gP_Br/Wa_–values and tested cumulatively for association with AD by Broad/WashU PGSA.

#### Follow-up BroWas/Baylor PGSA

Significant genes and polygenic components identified by Broad/WashU PGSA were tested in independent samples using BroWas/Baylor PGSA. For these follow-up analyses, CEN_TREM2_ variants were analyzed by logistic regression using combined Broad and WashU data, and BroWas-derived polygenic scores were tested for association with AD in the Baylor data.

### Optimal sequence kernel association testing (SKAT-O)

To determine if variants with minor allele counts less than 20 in polygenic components that showed significant association using PGSA are also associated with AD, we used SKAT-O. SKAT-O maximizes power by optimally combining burden test, which is typically used when most variants in the tested set are causal and their effects are in the same direction, with the non-burden sequence kernel association test, which is mostly used when a large fraction of variants are noncausal or direction of causal and noncausal variants are different directions^22,23^.

## Acknowledgements

We thank Shannon McDonnell for managing QC analysis of ADSP subjects and variants. We thank the patients and their families for their participation, without which these studies would not have been possible. This work was supported by National Institute on Aging [U01AG046139 to NE-T and SGY; RF AG051504;; R01AG061796 to NE-T].

For samples collected through the Sun Health Research Institute Brain and Body Donation Program of Sun City, Arizona and utilized in the brain expression studies: The Brain and Body Donation Program is supported by the National Institute of Neurological Disorders and Stroke (U24NS072026 National Brain and Tissue Resource for Parkinson’s Disease and Related Disorders), the National Institute on Aging (P30 AG19610 Arizona Alzheimer’s Disease Core Center), the Arizona Department of Health Services (contract 211002, Arizona Alzheimer’s Research Center), the Arizona Biomedical Research Commission (contracts 4001, 0011, 05-901, and 1001 to the Arizona Parkinson’s Disease Consortium), and the Michael J. Fox Foundation for Parkinson’s Research.

## Acknowledgment Statement for the ADSP

*The Alzheimer’s Disease Sequencing Project (ADSP) is comprised of two Alzheimer’s Disease (AD) genetics consortia and three National Human Genome Research Institute (NHGRI) funded Large Scale Sequencing and Analysis Centers (LSAC). The two AD genetics consortia are the Alzheimer’s Disease Genetics Consortium (ADGC) funded by NIA (U01 AG032984), and the Cohorts for Heart and Aging Research in Genomic Epidemiology (CHARGE) funded by NIA (R01 AG033193), the National Heart, Lung, and Blood Institute (NHLBI), other National Institute of Health (NIH) institutes and other foreign governmental and non-governmental organizations. The Discovery Phase analysis of sequence data is supported through UF1AG047133 (to Drs. Schellenberg, Farrer, Pericak-Vance, Mayeux, and Haines); U01AG049505 to Dr. Seshadri; U01AG049506 to Dr. Boerwinkle; U01AG049507 to Dr. Wijsman; and U01AG049508 to Dr. Goate and the Discovery Extension Phase analysis is supported through U01AG052411 to Dr. Goate, U01AG052410 to Dr. Pericak-Vance and U01 AG052409 to Drs. Seshadri and Fornage. Data generation and harmonization in the Follow-up Phases is supported by U54AG052427 (to Drs. Schellenberg and Wang).*

*The ADGC cohorts include: Adult Changes in Thought (ACT), the Alzheimer’s Disease Centers (ADC), the Chicago Health and Aging Project (CHAP), the Memory and Aging Project (MAP), Mayo Clinic (MAYO), Mayo Parkinson’s Disease controls, University of Miami, the Multi-Institutional Research in Alzheimer’s Genetic Epidemiology Study (MIRAGE), the National Cell Repository for Alzheimer’s Disease (NCRAD), the National Institute on Aging Late Onset Alzheimer’s Disease Family Study (NIA-LOAD), the Religious Orders Study (ROS), the Texas Alzheimer’s Research and Care Consortium (TARC), Vanderbilt University/Case Western Reserve University (VAN/CWRU), the Washington Heights-Inwood Columbia Aging Project (WHICAP) and the Washington University Sequencing Project (WUSP), the Columbia University Hispanic-Estudio Familiar de Influencia Genetica de Alzheimer (EFIGA), the University of Toronto (UT), and Genetic Differences (GD).*

*The CHARGE cohorts are supported in part by National Heart, Lung, and Blood Institute (NHLBI) infrastructure grant HL105756 (Psaty), RC2HL102419 (Boerwinkle) and the neurology working group is supported by the National Institute on Aging (NIA) R01 grant AG033193. The CHARGE cohorts participating in the ADSP include the following: Austrian Stroke Prevention Study (ASPS), ASPS-Family study, and the Prospective Dementia Registry-Austria (ASPS/PRODEM-Aus), the Atherosclerosis Risk in Communities (ARIC) Study, the Cardiovascular Health Study (CHS), the Erasmus Rucphen Family Study (ERF), the Framingham Heart Study (FHS), and the Rotterdam Study (RS). ASPS is funded by the Austrian Science Fond (FWF) grant number P20545-P05 and P13180 and the Medical University of Graz. The ASPS-Fam is funded by the Austrian Science Fund (FWF) project I904),the EU Joint Programme - Neurodegenerative Disease Research (JPND) in frame of the BRIDGET project (Austria, Ministry of Science) and the Medical University of Graz and the Steiermärkische Krankenanstalten Gesellschaft. PRODEM-Austria is supported by the Austrian Research Promotion agency (FFG) (Project No. 827462) and by the Austrian National Bank (Anniversary Fund, project 15435. ARIC research is carried out as a collaborative study supported by NHLBI contracts (HHSN268201100005C, HHSN268201100006C, HHSN268201100007C, HHSN268201100008C, HHSN268201100009C, HHSN268201100010C, HHSN268201100011C, and HHSN268201100012C). Neurocognitive data in ARIC is collected by U01 2U01HL096812, 2U01HL096814, 2U01HL096899, 2U01HL096902, 2U01HL096917 from the NIH (NHLBI, NINDS, NIA and NIDCD), and with previous brain MRI examinations funded by R01-HL70825 from the NHLBI. CHS research was supported by contracts HHSN268201200036C, HHSN268200800007C, N01HC55222, N01HC85079, N01HC85080, N01HC85081, N01HC85082, N01HC85083, N01HC85086, and grants U01HL080295 and U01HL130114 from the NHLBI with additional contribution from the National Institute of Neurological Disorders and Stroke (NINDS). Additional support was provided by R01AG023629, R01AG15928, and R01AG20098 from the NIA. FHS research is supported by NHLBI contracts N01-HC-25195 and HHSN268201500001I. This study was also supported by additional grants from the NIA (R01s AG054076, AG049607 and AG033040 and NINDS (R01 NS017950). The ERF study as a part of EUROSPAN (European Special Populations Research Network) was supported by European Commission FP6 STRP grant number 018947 (LSHG-CT-2006-01947) and also received funding from the European Community’s Seventh Framework Programme (FP7/2007-2013)/grant agreement HEALTH-F4-2007-201413 by the European Commission under the programme “Quality of Life and Management of the Living Resources” of 5th Framework Programme (no. QLG2-CT-2002-01254). High-throughput analysis of the ERF data was supported by a joint grant from the Netherlands Organization for Scientific Research and the Russian Foundation for Basic Research (NWO-RFBR 047.017.043). The Rotterdam Study is funded by Erasmus Medical Center and Erasmus University, Rotterdam, the Netherlands Organization for Health Research and Development (ZonMw), the Research Institute for Diseases in the Elderly (RIDE), the Ministry of Education, Culture and Science, the Ministry for Health, Welfare and Sports, the European Commission (DG XII), and the municipality of Rotterdam. Genetic data sets are also supported by the Netherlands Organization of Scientific Research NWO Investments (175.010.2005.011, 911-03-012), the Genetic Laboratory of the Department of Internal Medicine, Erasmus MC, the Research Institute for Diseases in the Elderly (014-93-015; RIDE2), and the Netherlands Genomics Initiative (NGI)/Netherlands Organization for Scientific Research (NWO) Netherlands Consortium for Healthy Aging (NCHA), project 050-060-810. All studies are grateful to their participants, faculty and staff. The content of these manuscripts is solely the responsibility of the authors and does not necessarily represent the official views of the National Institutes of Health or the U.S. Department of Health and Human Services.*

*The four LSACs are: the Human Genome Sequencing Center at the Baylor College of Medicine (U54 HG003273), the Broad Institute Genome Center (U54HG003067), The American Genome Center at the Uniformed Services University of the Health Sciences (U01AG057659), and the Washington University Genome Institute (U54HG003079)*.

*Biological samples and associated phenotypic data used in primary data analyses were stored at Study Investigators institutions, and at the National Cell Repository for Alzheimer’s Disease (NCRAD, U24AG021886) at Indiana University funded by NIA. Associated Phenotypic Data used in primary and secondary data analyses were provided by Study Investigators, the NIA funded Alzheimer’s Disease Centers (ADCs), and the National Alzheimer’s Coordinating Center (NACC, U01AG016976) and the National Institute on Aging Genetics of Alzheimer’s Disease Data Storage Site (NIAGADS, U24AG041689) at the University of Pennsylvania, funded by NIA, and at the Database for Genotypes and Phenotypes (dbGaP) funded by NIH. This research was supported in part by the Intramural Research Program of the National Institutes of health, National Library of Medicine. Contributors to the Genetic Analysis Data included Study Investigators on projects that were individually funded by NIA, and other NIH institutes, and by private U.S. organizations, or foreign governmental or nongovernmental organizations.*

## Author information

J.S.R and S.G.Y. conceived the project and performed all of the PGSA and SKAT-O analyses. J.M.B and G.D.J collaborated closely with J.S.R and S.G.Y in the early stages of the PGSA and SKAT-O analyses. M.A, X.W., and N.E-T performed WGCNA. All authors were involved in the design and/or execution of single variant analysis and QC. J.S.R. and S.G.Y prepared the first draft of the manuscript. All authors contributed to the final manuscript.

## Supplementary Figures

**Supplementary Figure 1.**
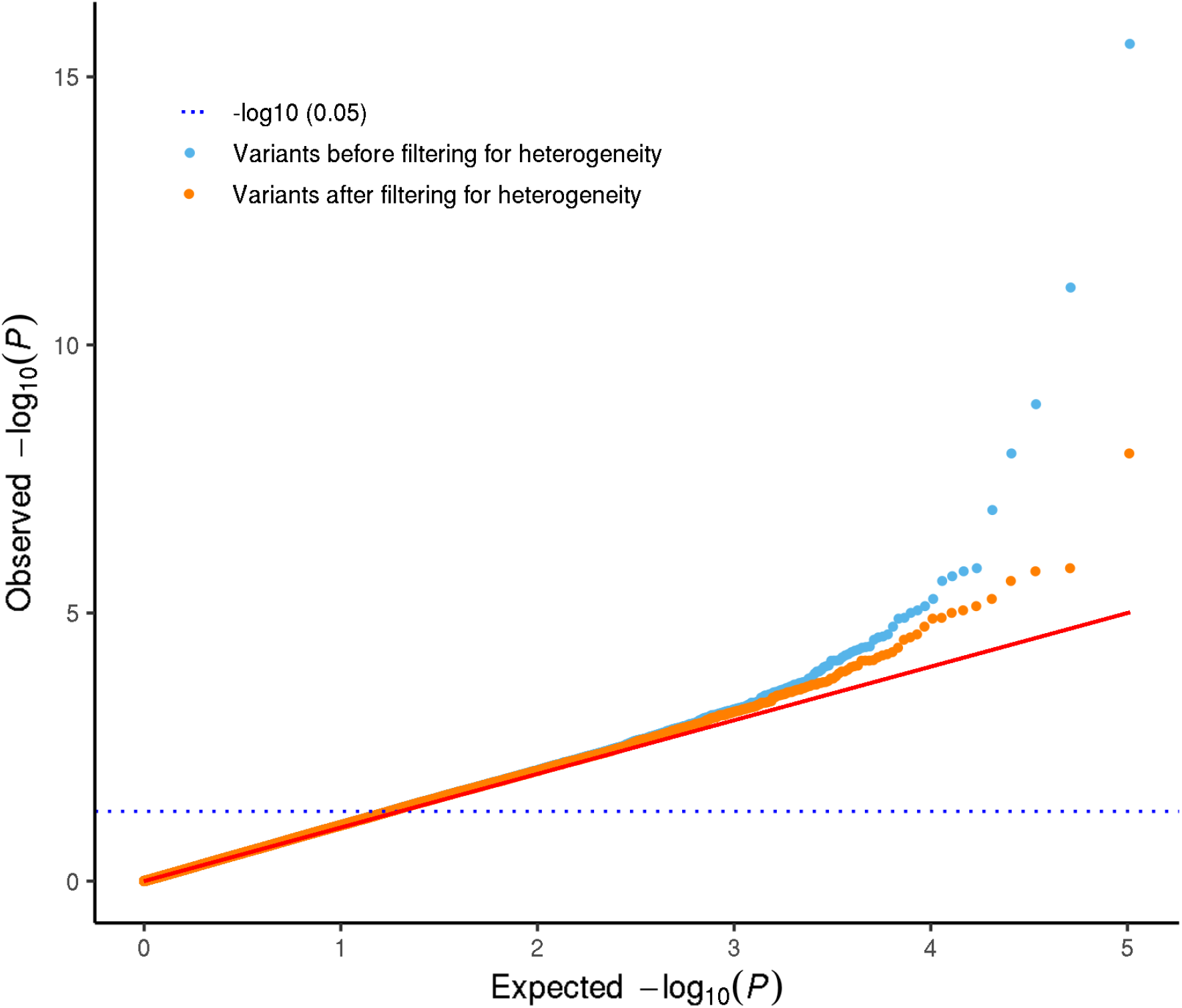
Q-Q plots of P_ADSP_-values of ADSP WES variants before and after filtering for heterogeneity. Using all 9904 post-QC ADSP subjects, all post-QC variants with a minor allele count ≥ 20 were analyzed by logistic regression using an additive model with sex*, APOE* ε4 dose, *APOE* ε2 dose, LSACs, and three principal component vectors as covariates. As a final QC measure, multinomial regression was performed to assess heterogeneity in minor allele frequency across the three LSACs. Q-Q plots of P_ADSP_ values are shown before (light blue points) and after (orange points) removing, 677 variants (0.66%) with study-wide P_LSAC_ values ≤ 4.86E-07.

**Supplementary Figure 2.**
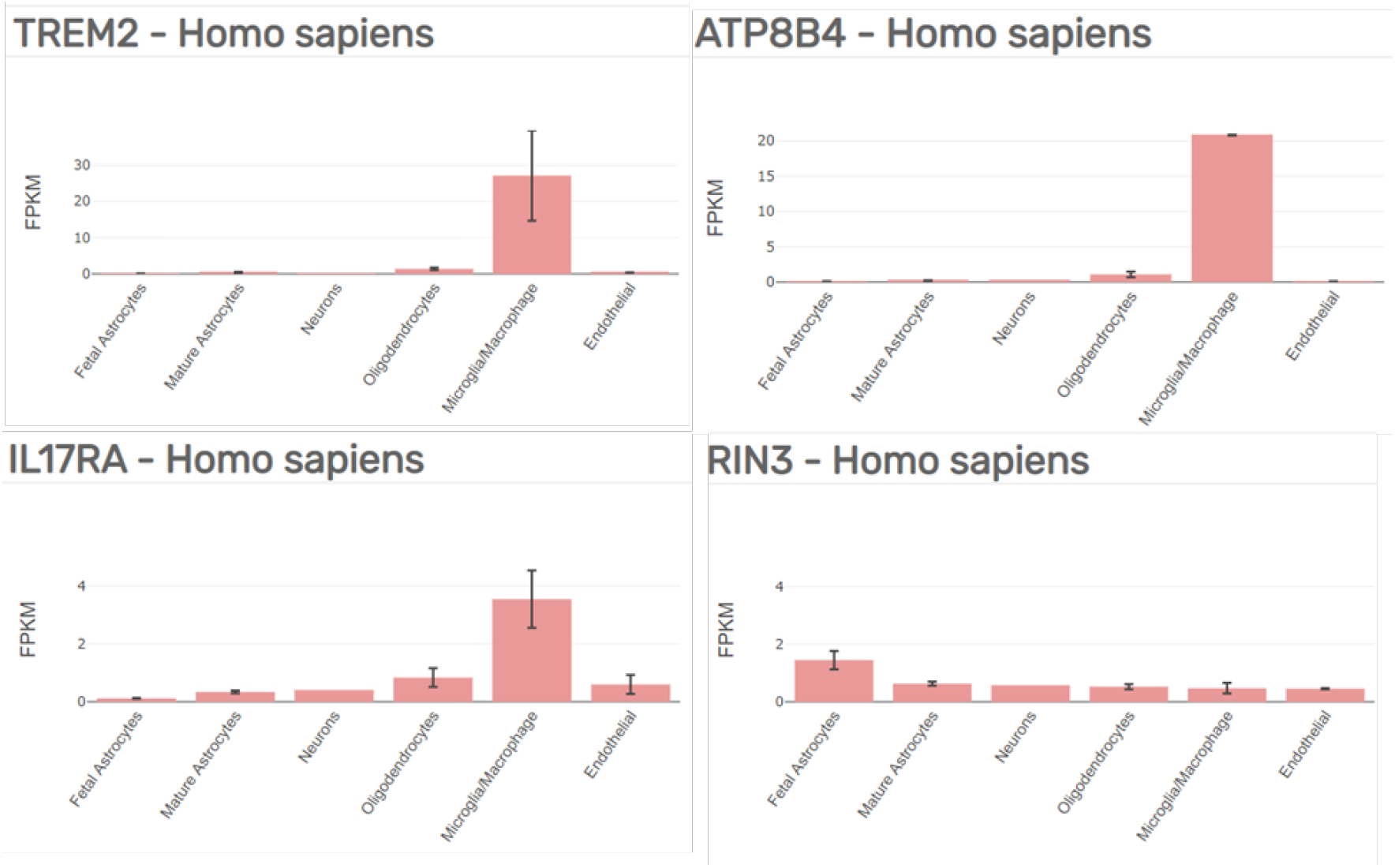
Gene expression in human brain cells. (Source: Zhang, et al, 2016, https://www.brainrnaseq.org/). Gene expression values for the five novel (*TREM2, ATP8B4, IL17RA and RIN3*) and two known AD genes (*TREM2* and *RIN3*) as observed in human CNS cell types.

## References

1. Holtzman, D.M., Morris, J.C. & Goate, A.M. Alzheimer’s disease: the challenge of the second century. Sci Transl Med 3, 77sr1 (2011).

2. Jack, C.R., Jr. et al. Hypothetical model of dynamic biomarkers of the Alzheimer’s pathological cascade. Lancet Neurol 9, 119–28 (2010).

3. Ransohoff, R.M. How neuroinflammation contributes to neurodegeneration. Science 353, 777–83 (2016).

4. Shi, Y. & Holtzman, D.M. Interplay between innate immunity and Alzheimer disease: APOE and TREM2 in the spotlight. Nat Rev Immunol 18, 759–772 (2018).

5. Jonsson, T. et al. Variant of TREM2 associated with the risk of Alzheimer’s disease. N Engl J Med 368, 107–16 (2013).

6. Guerreiro, R. et al. TREM2 variants in Alzheimer’s disease. N Engl J Med 368, 117–27 (2013).

7. Zhang, Y. et al. Purification and Characterization of Progenitor and Mature Human Astrocytes Reveals Transcriptional and Functional Differences with Mouse. Neuron 89, 37–53 (2016).

8. Ma, L. et al. Expression and processing analyses of wild type and p.R47H TREM2 variant in Alzheimer’s disease brains. Mol Neurodegener 11, 72 (2016).

9. Conway, O.J. et al. ABI3 and PLCG2 missense variants as risk factors for neurodegenerative diseases in Caucasians and African Americans. Mol Neurodegener 13, 53 (2018).

10. Sims, R. et al. Rare coding variants in PLCG2, ABI3, and TREM2 implicate microglial-mediated innate immunity in Alzheimer’s disease. Nat Genet 49, 1373–1384 (2017).

11. Ulrich, J.D., Ulland, T.K., Colonna, M. & Holtzman, D.M. Elucidating the Role of TREM2 in Alzheimer’s Disease. Neuron 94, 237–248 (2017).

12. Gratuze, M. et al. Impact of TREM2R47H variant on tau pathology-induced gliosis and neurodegeneration. J Clin Invest (2020).

13. Sayed, F.A. et al. Differential effects of partial and complete loss of TREM2 on microglial injury response and tauopathy. Proc Natl Acad Sci U S A 115, 10172–10177 (2018).

14. Leyns, C.E.G. et al. TREM2 deficiency attenuates neuroinflammation and protects against neurodegeneration in a mouse model of tauopathy. Proc Natl Acad Sci U S A 114, 11524–11529 (2017).

15. Matarin, M. et al. A genome-wide gene-expression analysis and database in transgenic mice during development of amyloid or tau pathology. Cell Rep 10, 633–44 (2015).

16. Zhang, B. et al. Integrated systems approach identifies genetic nodes and networks in late-onset Alzheimer’s disease. Cell 153, 707–20 (2013).

17. Langfelder, P. & Horvath, S. WGCNA: an R package for weighted correlation network analysis. BMC Bioinformatics 9, 559 (2008).

18. Allen, M. et al. Human whole genome genotype and transcriptome data for Alzheimer’s and other neurodegenerative diseases. Sci Data 3, 160089 (2016).

19. Beecham, G.W. et al. The Alzheimer’s Disease Sequencing Project: Study design and sample selection. Neurol Genet 3, e194 (2017).

20. Lambert, J.C. et al. Meta-analysis of 74,046 individuals identifies 11 new susceptibility loci for Alzheimer’s disease. Nat Genet 45, 1452–8 (2013).

21. Rentzsch, P., Witten, D., Cooper, G.M., Shendure, J. & Kircher, M. CADD: predicting the deleteriousness of variants throughout the human genome. Nucleic Acids Res 47, D886–D894 (2019).

22. Lee, S. et al. Optimal unified approach for rare-variant association testing with application to small-sample case-control whole-exome sequencing studies. Am J Hum Genet 91, 224–37 (2012).

23. Ionita-Laza, I., Lee, S., Makarov, V., Buxbaum, J.D. & Lin, X. Sequence kernel association tests for the combined effect of rare and common variants. Am J Hum Genet 92, 841–53 (2013).

24. Kunkle, B.W. et al. Genetic meta-analysis of diagnosed Alzheimer’s disease identifies new risk loci and implicates Abeta, tau, immunity and lipid processing. Nat Genet 51, 414–430 (2019).

25. Jansen, I.E. et al. Genome-wide meta-analysis identifies new loci and functional pathways influencing Alzheimer’s disease risk. Nat Genet 51, 404–413 (2019).

26. Holstege, H. et al. Exome sequencing identifies novel AD-associated genes. medRxiv, 2020.07.22.20159251 (2020).

27. Wang, M. et al. The Mount Sinai cohort of large-scale genomic, transcriptomic and proteomic data in Alzheimer’s disease. Sci Data 5, 180185 (2018).

28. St John-Williams, L. et al. Targeted metabolomics and medication classification data from participants in the ADNI1 cohort. Sci Data 4, 170140 (2017).

29. Van der Auwera, G.A. et al. From FastQ data to high confidence variant calls: the Genome Analysis Toolkit best practices pipeline. Curr Protoc Bioinformatics 43, 11 10 1–33 (2013).

30. Quinlan, A.R. & Hall, I.M. BEDTools: a flexible suite of utilities for comparing genomic features. Bioinformatics 26, 841–2 (2010).

31. Chang, C.C. et al. Second-generation PLINK: rising to the challenge of larger and richer datasets. Gigascience 4, 7 (2015).

32. Jun, G. et al. Detecting and estimating contamination of human DNA samples in sequencing and array-based genotype data. Am J Hum Genet 91, 839–48 (2012).

33. Manichaikul, A. et al. Robust relationship inference in genome-wide association studies. Bioinformatics 26, 2867–73 (2010).

34. Patterson, N., Price, A.L. & Reich, D. Population structure and eigenanalysis. PLoS Genet 2, e190 (2006).

35. Wang, K., Li, M. & Hakonarson, H. ANNOVAR: functional annotation of genetic variants from high-throughput sequencing data. Nucleic Acids Res 38, e164 (2010).

36. Langfelder, P. & Horvath, S. WGCNA: an R package for weighted correlation network analysis. BMC Bioinformatics 9, 559 (2008).

37. Allen, M. et al. Conserved brain myelination networks are altered in Alzheimer’s and other neurodegenerative diseases. Alzheimer’s & Dementia (2017).

38. Kalari, K.R. et al. MAP-RSeq: Mayo Analysis Pipeline for RNA sequencing. BMC Bioinformatics 15, 224 (2014).

39. Hansen, K.D., Irizarry, R.A. & Wu, Z. Removing technical variability in RNA-seq data using conditional quantile normalization. Biostatistics 13, 204–216 (2012).

